# Year-long transcriptome survey highlights a resource allocation strategy at the late growth stage in a monocarpic plant

**DOI:** 10.1101/2025.10.14.682294

**Authors:** Yasuyuki Nomura, Atsushi J. Nagano

## Abstract

Day length and temperature drastically changes along the annual cycle in temperate zone. Temperate plants are thought to adapt to the annual cycle by various physiological responses in appropriate season. In the previous study, the annual transcriptome dynamics of *Arabidopsis halleri*, which is a polycarpic sister species of *A. thaliana*, was obtained in natural habitats and annual environments simulated by growth chambers. Here, we performed year-long transcriptome analysis of *A. thaliana* (Col-0 and FRI^Sf2^) under annual environments simulated by growth chambers. Comparing gene expressions in FRI^Sf2^ and *A. helleri,* most genes had similar annual expression pattens. However, in February-March conditions, 2,787 genes, containing *LHCB* and *TGG1*, exhibited significantly lower expressions in FRI^Sf2^ than those in *A. helleri. LHCB* and *TGG1* were not downregulated in cold treatments for a month. These results suggest that factors other than temperature, such as age, are involved in the down-regulations of *LHCB* and *TGG1*. Resource translocation to seeds occurs at the end of the growing season in annual plants. The down-regulation of photosynthesis-related proteins and defense-related proteins, known to be highly abundant in leaves, is possibly an adaptive strategy of monocarpic *A. thaliana*.

## INTRODUCTION

Most plants grow for several months or longer, and during this time, they are exposed to a variety of environmental changes. Typical examples include changes in light conditions, temperature, and humidity over a few hours, as well as changes in day length over several months. Plants respond to these environmental fluctuations, which enable them to undergo germination, leaf spreading, flowering, defoliation, and dormancy at the appropriate time (Polgar & Primack, 2011; Hepworth & Dean, 2015; Kudoh, 2016; Penfield & MacGregor, 2017). In addition to these visible seasonal events, physiological states are also altered by environmental cues (Keskitalo *et al*., 2005). Elucidating the gene expression regulation underlying these events is essential for understanding plant life history. Not only extrinsic factors like temperature and day length, but also intrinsic factors such as the circadian clock, nutritional conditions, and age, regulate the gene expression that controls the timing of life history events (Aikawa *et al*., 2010; Miyazaki *et al*., 2014; Satake *et al*., 2019). For example, floral initiation in *Arabidopsis thaliana* (*Ath*) occurs after long-term cultivation (Lee *et al*., 1993; Lee & Amasino, 1995), suggesting that age can also induce flowering.

A previous study obtained transcriptomes of *A. halleri* subsp. *gemmifera* (*Ahg*), a wild perennial species closely related to *Ath*, weekly in natural habitats for two years and biweekly in a growth chamber simulating annual pattern (Nagano *et al*., 2019). This revealed that oscillated genes along the annual cycle can be explained by temperature. The expression levels of the *FLC* (*FLOWERING LOCUS C*) gene, one of the seasonally-oscillated genes, could be explained by temperatures over the preceding six weeks, indicating a long-term response spanning a month or more (Aikawa *et al*., 2010). In contrast, when *Ath* was cultured for one, three, or seven days under temperature and day-length conditions corresponding to each month, some genes mimicked the monthly gene expression of *Ahg*, while others did not (Kurita *et al*., 2021). From a comparison of these data, it was not possible to distinguish whether the inability to mimic gene expression was due to species differences or the short cultivation period. Therefore, it is necessary to compare data from long-term cultivation of *Ahg* with data from long-term cultivation of *Ath*.

By comparing monocarpic (annual) and polycarpic (perennial) plants, we expect clear differences in gene expression levels, especially during the flowering period. These differences are not limited to flowering responses but also extend to resource allocation. Specifically, annual plants allocate the majority of their internal resources to seed production, whereas perennial plants distribute only a portion of their resources to seeds. In this study, we cultivated *Ath* (Col-0 and FRI^Sf2^) for ten months in growth chambers that simulated the annual environmental pattern to reveal gene expression variation over several months to a year. From these experiments, we found *Ath*-specific down-regulations of photosynthesis- and defence-related genes that may be involved in the differences in the life history strategies of *Ahg* and *Ath*.

## MATERIALS AND METHODS

### Plant materials and common settings of growth chamber

As stratification and pre-cultivation, seeds of two *Ath* lines, Col-0 (CS70000) and FRI^Sf2^ (Lee *et al*., 1993), were sown in each pot of 7 cm diameter filled with vermiculite with liquid fertilizer (N;1.8 mg, P; 3.0 mg, K; 1.5 mg per pot, HYPONeX, HYPONeX Japan, Osaka, Japan), and placed under 10°C with dark for one day. Then, the pots were transferred to a condition under 20°C with 8 h light / 16 h dark cycle for 35 days (Tables S1-S5). Photosynthetic photon flux density was set at about 83 μ mol s^−1^ m^−2^ at the soil surface.

### Growth chamber experiments simulating annual cycle

The GC experiment simulating annual cycle was conducted in two periods for Col-0, from July to September and from January to April, and in one period for FRI^Sf2^, from July to April. 162 plants of Col-0 were divided and transferred into two growth chambers (HCLP-880PFD-3-4LS, NK System, Osaka, Japan) set for natural and in-phase conditions (Figure 1A, Table S1). Col-0 plants were grown under each condition for about four months (Table S2). Because Col-0 flowered and died at two months after from the start of each condition, the second series using only Col-0 started after 154 days later from the start of the first series. 162 Col-0 plants were divided and transferred into two growth chambers as same as the first series. The second series plants were grown under each condition for about five months (Table S3). 162 plants of FRI^Sf2^ were also divided and transferred into two growth chambers set for natural and in-phase conditions (Figure 1A, Table S1). FRI^Sf2^ plants were grown under each condition for ten months (Table S2).

**Figure 1.**
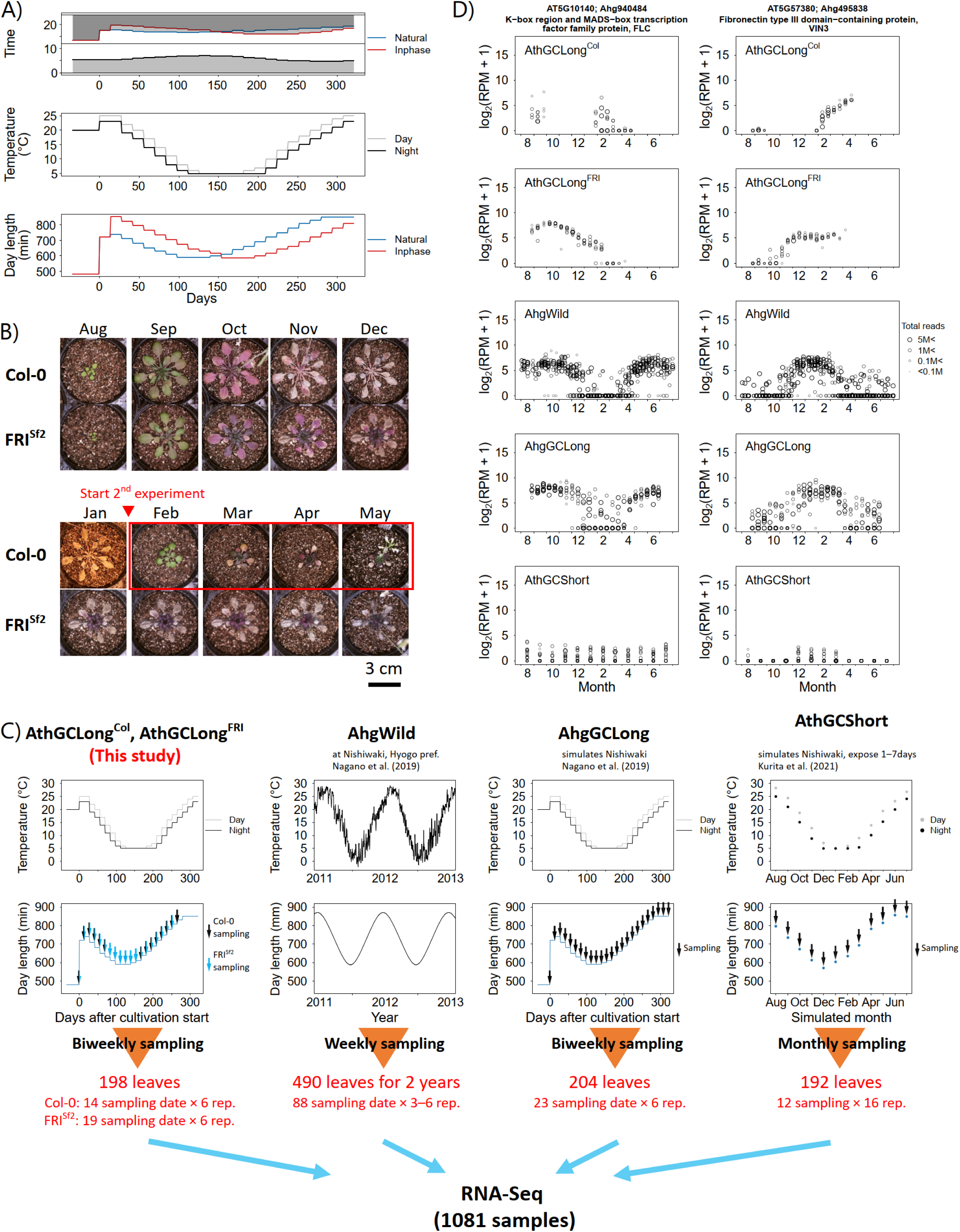
Design of the experiments and an example of the results. (A) Day length and temperature settings in this study. Shaded areas in the upper panel indicate dark period. The horizontal black line indicates the time of sampling, 12:00. (B) Col-0 and FRI^Sf2^ in each month. Col-0 died in January, so a second experiment was conducted starting in January. (C) Temperature, day length and sampling design in this study and related studies for comparison (Nagano *et al*., 2019; Kurita *et al*., 2021). (D) Annual expression pattern of *FLC* and *VIN3* in Col-0 and FRI^Sf2^.

Day length settings and temperature settings of the growth chambers which were set with reference to the previous study (Nagano *et al*., 2019). They were determined on the basis of two week-averaged values of theoretical sunrise and sunset times (National Astronomical Observatory of Japan) and air temperature records (Japan Meteorological Agency), respectively, at Nishiwaki, Japan (34° 59’ 36’’ 240 N, 134° 58’ 09’ 120 E). The number of flowers and the number of fruits longer than 5 mm produced during the experiment was measured at the end of the experiment. Leaves were sampled from 36, 48 and 114 plants of Col-0 in first (hereafter AthGCLong^Col1^), second series (hereafter AthGCLong^Col2^) and FRI^Sf2^ (hereafter AthGCLong^FRI^), respectively, and every two weeks by collecting an uppermost full-enrolled leaf at 14:00 (Table S1-S3). The leaves were immediately frozen in liquid nitrogen and stored at −80 °C until RNA extraction for RNA-Seq analysis.

### Growth chamber experiments exposing cold

In the GC experiment exposing cold, 64 plants of FRI^Sf2^ were divided and transferred into two growth chambers set for control and cold conditions (Table S4). For the control condition, the temperature was kept at 20°C for three months. For the cold condition, the temperature was set at 20°C for one month of pre-cultivation, 4°C for the next month, and 20°C for the last month. FRI^Sf2^ plants were grown under each condition for about three months (Table S5). All leaves were weekly sampled at 14:00 (Table S4). The leaves were immediately frozen in liquid nitrogen and stored at −80 °C until RNA extraction for RNA-Seq analysis.

### RNA-Seq experiment

Leaves were crushed with two or three zirconia beads, using the TissueLyser II (QIAGEN, MD, USA) with the pre-chilled adapters at −80 °C. Total RNA was extracted from leaves using Maxwell 16 LEV Plant RNA Kit (Promega, WI, USA). The amount of RNA was measured using Quant-iT RNA Assay Kit broad range (Thermo Fisher Scientific, Waltham, MA, USA) and Tecan plate reader Infinite 200 PRO (Tecan, Männedorf, Switzerland). For RNA-Seq library preparation, 400 or 500 ng of total RNA per sample was used. The libraries were prepared using Lasy-Seq v1.1 protocol (Kamitani *et al*., 2019) (https://sites.google.com/view/lasy-seq/). The quality of the libraries was assessed by the Bioanalyzer (Agilent Technologies). The libraries were sequenced by the paired-end 150 bases with the HiSeq X (Illumina).

Illumina adapter removal and low-quality filtering (Q < 20) of RNA-Seq data were performed using the Trimmomatic-0.33 (Bolger *et al*., 2014). Then, preprocessed reads were mapped on transcript sequences in TAIR 10 with Bowtie (v1.1.1) (Langmead *et al*., 2009) and quantified using RSEM-1.2.21(Li & Dewey, 2011). Conversion of output from RSEM to read per million (RPM) was performed using R (version 4) (R Core Team, 2021) in the same manner as in the previous study (Kamitani *et al*., 2016).

### Data sets from previous studies

RNA-Seq data of *Ahg* were obtained from previous study (Nagano *et al*., 2019). *Ahg* leaves in nature (hereafter AhgWild) were collected weekly from the habitats in middle part of Japan (the Omoide River, Taka-cho, 35° 06’ N, 134° 55’ E) (Figure 1). Half-sib *Ahg* leaves in the GC experiment (hereafter AhgGCLong) were also collected biweekly under the same condition with GC experiment simulation annual cycle mentioned above (Figure 1).

*Ath* Col-0 leaves were collected after 1, 3, and 7 days of exposure to conditions that mimicked the monthly average of temperature and day length from January to December in Nishiwaki (34° 59’ 36’’ 240 N, 134° 58’ 09’ 120 E) (hereafter AthGCShort) (Kurita *et al*., 2021).

RNA-Seq data of *Ath* and *Ah*g were analyzed as follows (Figure S2). Samples with fewer than 10^5.5^ reads were excluded from the analysis. Sample attributes used in the analysis were shown in Table S1. 22,845 *Ahg* genes was annotated as orthologs of *Ath* genes by reciprocal blast. Genes with log_2_(RPM+1) > 2 on average throughout all samples (*Ath* and *Ahg*) were defined as expressed genes. Among the expressed genes, genes with a standard deviation of log_2_(RPM+1) > 1 in any of the expression profiles were defined as variable genes. Subsequent analysis was performed using the variable genes.

### Statistical analysis for RNA-Seq data

After log_2_(RPM+1) transformation of RPM, averages were calculated monthly for each data sets (AhgWild, AhgGCLong, AthGCShort, AthGCLong^Col^ and AthGCLong^FRI^). The Pearson’s correlation coefficient for log_2_(RPM+1) averaged by month was calculated. Brunner-Munzel test was performed to compare means for the correlation coefficients between AthGCShort-AhgWild and AthGCShort-AhgGCLong.

Differentially expressed gene (DEG) analysis was conducted using edgeR package (Robinson *et al*., 2010). Between AthWild and AthGCLong^FRI^, DEG detection was conducted with false discovery rate (FDR) < 0.05 from October to November and from February to March. Gene ontology (GO) enrichment analysis was performed on genes that were not DEGs in October-November but were DEGs in February-March using the GO.db package (Carlson, 2018), as described in the previous study (Nagano *et al*., 2019). The FDR was controlled using Benjamini and Hochberg’s method (Benjamini & Hochberg, 1995).

### Statistical analysis for other data

Generalized linear mixed models (GLMMs) were used to examine the associations among the variables. We constructed a GLMM with the treatment conditions (Natural and In-phase) as the explanatory variable. The response variables were the bolting date, the number of flowers, and the number of fruits. We modelled the bolting date using a Gaussian distribution with an identity link function, while the number of flowers and fruits were modelled with a Poisson distribution and a log link function. The treatment conditions were treated as a fixed effect, and the position of the plants was included as a random effect. We also conducted a GLMM with the treatment conditions and the bolting date as the explanatory factors. The response variable was the number of fruits. We modelled the number of fruits using Poisson distribution and a log link function. The treatment conditions, the bolting date and their interaction were treated as fixed effects. The position of plants was treated as a random effect. A likelihood ratio test was used to test the significance level of the fixed effects.

All analyses were performed using R (version 4) (R Core Team, 2021). GLMMs were constructed using the package “glmmTMB” (Brooks *et al*., 2017). The likelihood ratio test was performed using the ANOVA function in the “car” package (Fox & Weisberg, 2011). Tukey-Krammer test was performed using the package “multcomp” (Hothorn *et al*., 2008).

## RESULTS

### Transcriptomes under real and simulated annual environments

To clarify long term trend of gene expression of *Ath* in simulated conditions, Col-0 and FRI^Sf2^ plants were grown under conditions from August to April, and the uppermost fully-expanded leaves were sampled biweekly (Figure 1A). Col-0 was bolted in October and died and the second experiment was started in February (Figures 1B, S1). FRI^Sf2^ continued to grow until April (Figure 1B). Finally, 36 leaves were sampled for Col-0 in the first experiment (AthGCLong^Col1^), 48 leaves were for Col-0 in the second experiment (AthGCLong^Col2^), and 114 leaves were for FRI^Sf2^ (AthGCLong^FRI^) (Table S1). These samples were analyzed by RNA-Seq and 789 M reads in total were obtained. After quality filtering, 141 samples were analyzed (Figure S2). Homologs of reciprocal top hits of *Ath* and *Ahg* were included in the analysis. Among these orthologs, we analyzed 12,141 genes whose expression were observed (Figure S2). Previous studies of *Ahg* (Nagano *et al*., 2019) showed a difference in the number of flowers between natural and in-phase conditions. However, the difference was not found in *Ath* of this study (Figure S1). Since there was little difference in gene expression, we did not distinguish between the natural and in-phase conditions in subsequent analyses.

To compare gene expression variation in annual environments, we employed three data sets in previous studies (Nagano *et al*., 2019; Kurita *et al*., 2021) (Figure 1C). The first data set was obtained by collecting *Ahg* weekly for two years in natural habitats (AhgWild, 490 samples). The second data set was obtained by collecting *Ahg* biweekly for one year in growth chamber that simulated an annual cycle (AhgGCLong, 204 samples). The third data set was obtained by exposing Col-0 to the simulated environment of each month for 1, 3, or 7 days (hereafter, AthGCShort, 189 samples).

We confirmed a clear annual expression pattern of *FLC*. The expression of *FLC* peaked at the end of September in AthGCLong^FRI^ (Figure 1D). This result is reasonable because *FLC*, which is involved in the vernalization requirement of *Ath*, is known to be down-regulated under low temperatures (Michaels & Amasino, 1999; Sheldon *et al*., 1999, 2000). In AhgWild and AhgGCLong, *FLC* also peaked at the end of September, as in AthGCLong^FRI^, and its expression decreased during the winter season (Figure 1D). For *FLC*, AthGCLong^FRI^ grown for several months was more similar to the annual expression profile of AhgWild and AhgGCLong than AthGCShort. *VIN3* had also a clear annual expression profile. As the *FLC* gene expression was down-regulated, the *VIN3* expression was gradually up-regulated, and reached saturation in December in AthGCLong^FRI^, AhgWild and AhgGCLong. *FLC* and *VIN3* expressions were always low in AthGCShort.

### FRI^Sf2^ in late stage has different transcriptome dynamics from *Ahg*

We compared transcriptome dynamics under various real/simulated annual conditions (Figures 2). Pearson correlation coefficients were calculated for the average expression levels in each month of each dataset on all pairs (AhgWild, AhgGCLong, AthGCShort, AthGCLong^Col^ and AthGCLong^FRI^) (Figures 2A, S3). Sample sizes for each month of each dataset ranged from 19 to 50 for AhgWild, 18 to 26 for AhgGCLong, 14 to 19 for AthGCShort, 4 to 11 for AthGCLong^Col^, and 4 to 18 for AthGCLong^FRI^. AthGCLong^FRI^ had similar transcriptomic profiles with AhgWild and AthGCShort in the fall, but the similarity decreased as the season moved into spring, suggesting that AthGCLong^FRI^ in spring is in different physiological status from other transcriptomic profiles (Figure 2). In addition, despite the similarity of conditions of AthGCShort and AhgGCLong, the transcriptomic profile of AthGCShort was significantly more similar to AhgWild than that for AhgGCLong (Brunner-Munzel test, test statistic = 64.5, *P* = 1.56×10^−125^) (Figure 2A).

**Figure 2.**
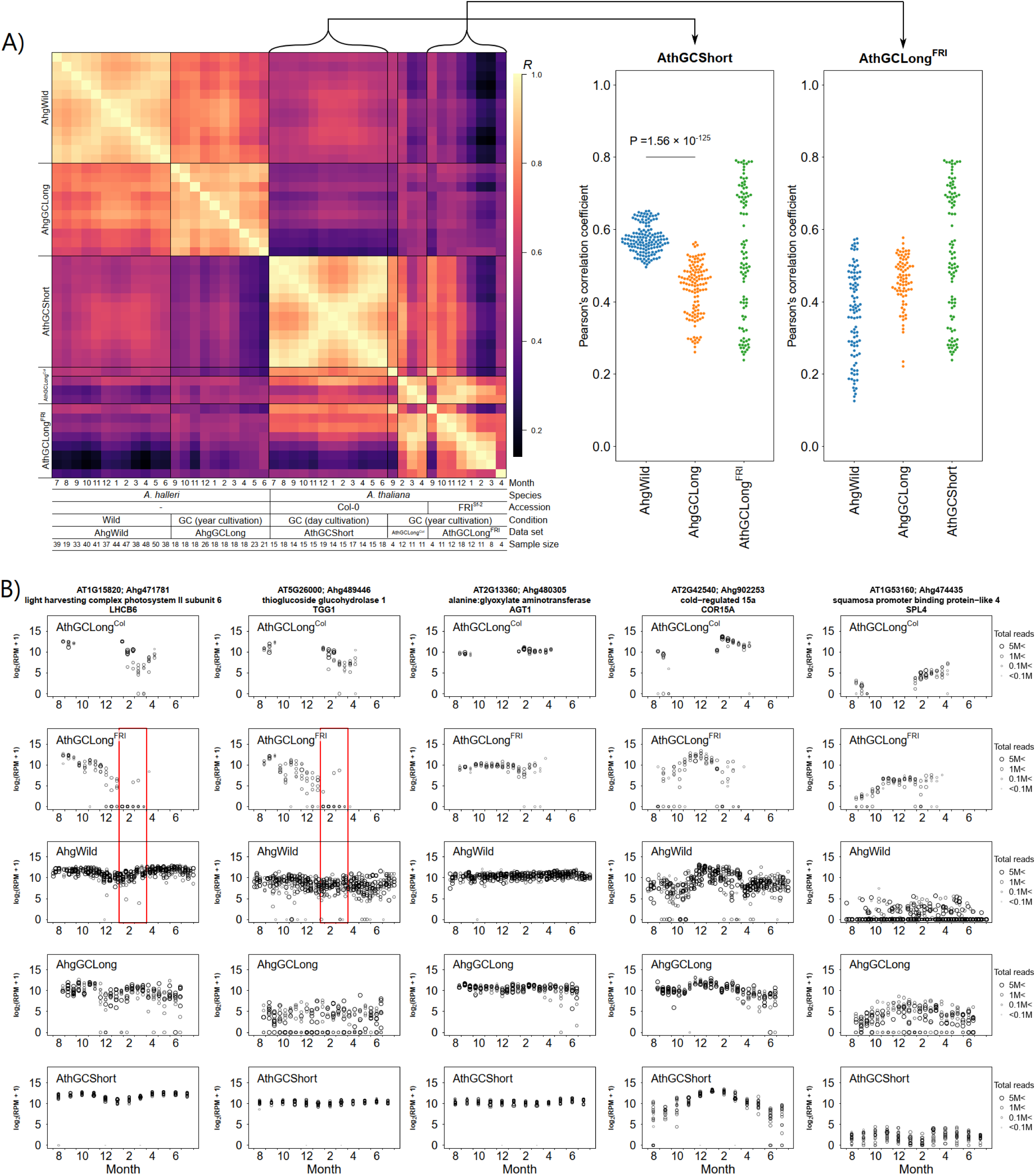
Annual transcriptome dynamics among studies. (A) Pearson correlations (*R*) between the monthly average of transcriptome profiles in the left panel. Correlation coefficients were calculated using the monthly average of log-transformed RPMs for the variable genes (n = 12,141 genes). Pearson correlation coefficients among AthGCLong^FRI^, AthGCShort and other data sets in the right panel. Pearson correlation coefficients of AthGCShort-AhgWild was significantly different from that of AthGCShort-AhgGCLong (Brunner-Munzel test). (B) Annual expression pattern of *LHCB6*, *TGG1*, *AGT*, *COR15A* and *SPL4*.

To clarify what made the unique transcriptomic profile, we looked for genes with unique expression status in AthGCLong^FRI^ in the spring. Differentially expressed genes (DEGs) between AthGCLong^FRI^ and AhgWild were tested in two time period: in Oct-Nov and in Feb-Mar. (Figure 3A). There were 1,367 genes that were not DEGs in October-November but were DEGs in February-March (Figure 3B). Gene ontology (GO) enrichment analysis using the down-regulated genes showed a significant enrichment of genes annotated as “photosynthesis”, “chloroplast organization” and “response to light stimulus” terms (Table S6). On the other hand, genes annotated with “leaf senescence”, “aging”, “defense response”, “transport” and “autophagy” were significantly enriched in up-regulated genes (Table S6).

**Figure 3.**
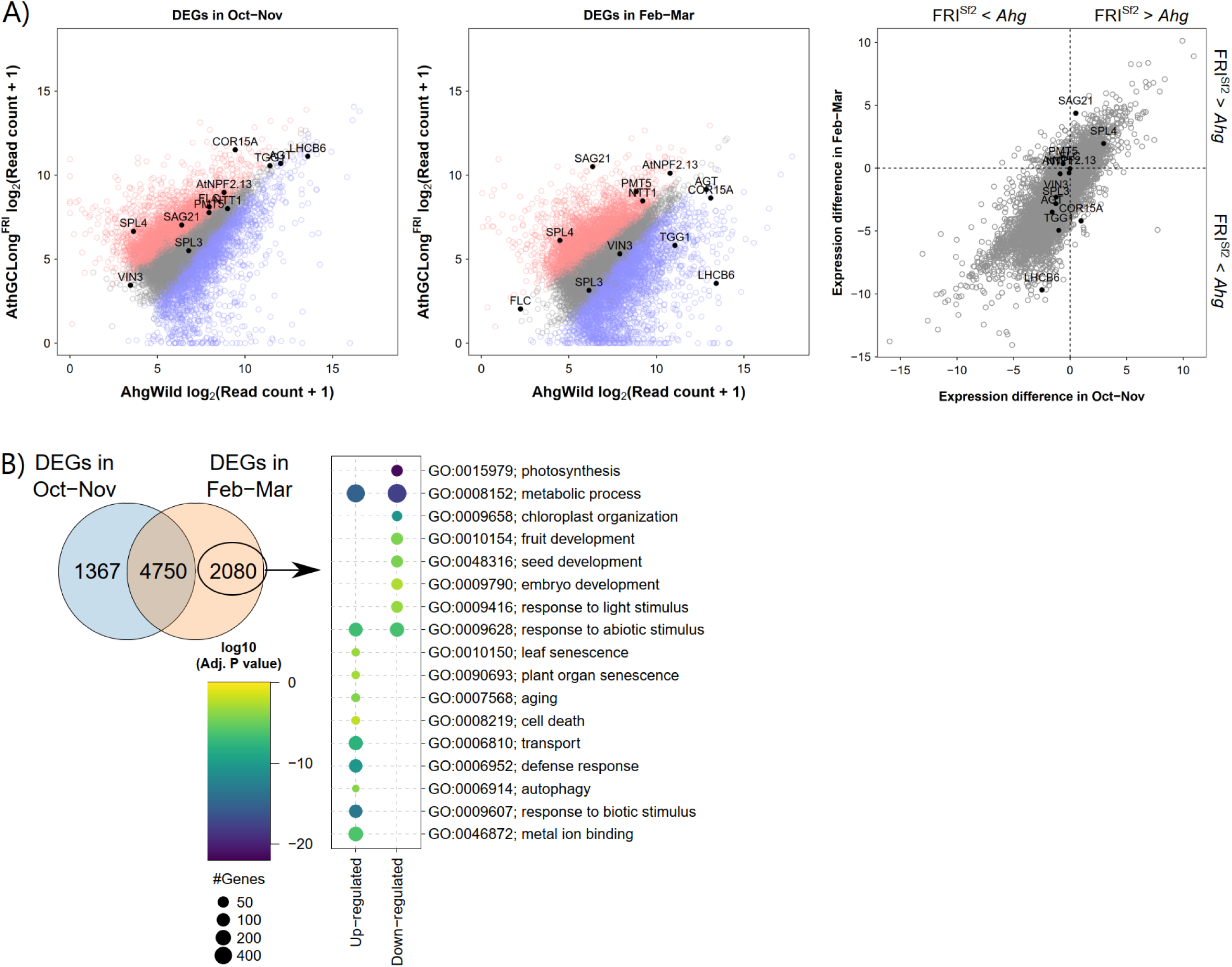
DEGs between *Ath* and *Ahg* in fall (October**–**November) and spring (February**–**March). (A) The left panel shows DEGs between AthGCLong^FRI^ and AhgWild in October– November. The centre panel shows DEGs between AthGCLong^FRI^ and AhgWild in February– March. The right panel shows the difference of gene expression between AthGCLong^FRI^ and AhgWild in fall (October–November) and spring (February–March). (B) The left panel shows Venn diagram of DEGs between AthGCLong^FRI^ and AhgWild. The right panel shows the result of GO term enrichment analysis of DEGs only in spring (February–March), not in fall (October–November).

In February-March, *LHCB6* and *TGG1* expression was significantly lower in AthGCLong^FRI^ than in AhgWild (adjusted *P* value = 1.0×10^−30^ and 6.4×10^−7^, respectively). For photosynthesis-related genes such as *LHCB6* and glucosinolate metabolism-related genes such as *TGG1*, expression gradually declined after December in AthGCLong^FRI^ and was barely expressed in April (Figure 2B, red box). This pattern of expression was not observed in AhgWild, AhgGCLong, and AthGCShort. Annual patterns of most genes in *Ath* were similar to that in *Ahg*. As an example of gene with annual cycle, *COR15a* with had a similar expression pattern in *Ahg* and *Ath*, with peak expression from November to February. As an example of less variable gene, *AGT* was always maintained at a constant expression level in *Ahg* and *Ath*. From these results, the unique transcriptome status of AthGCLong^FRI^ in spring is characterized from that photosynthesis- and glucosinolate metabolism-related genes remain down-regulated.

### Age, not cold, may play a role in decreased expression of *LHCB* and *TGG1*

To investigate whether the decrease in gene expression of *LHCB6* and *TGG1* during the late growing season was caused by low temperature or by plant age, FRI^Sf2^ plants were grown under two distinct conditions (Figure 4A). After a four-week precultivation period at a constant 20°C, the plants were divided into a group maintained at 20°C (control group) and a group exposed to 5°C (cold group) and cultivated for 4 weeks. After the cold treatment, the plants in the cold group were returned to a constant 20°C condition and cultivated for an additional 4 weeks. All leaves were sampled weekly from both groups. A total of 54 samples were collected and analyzed using RNA-Seq, yielding a total of 584 million read-pairs.

**Figure 4.**
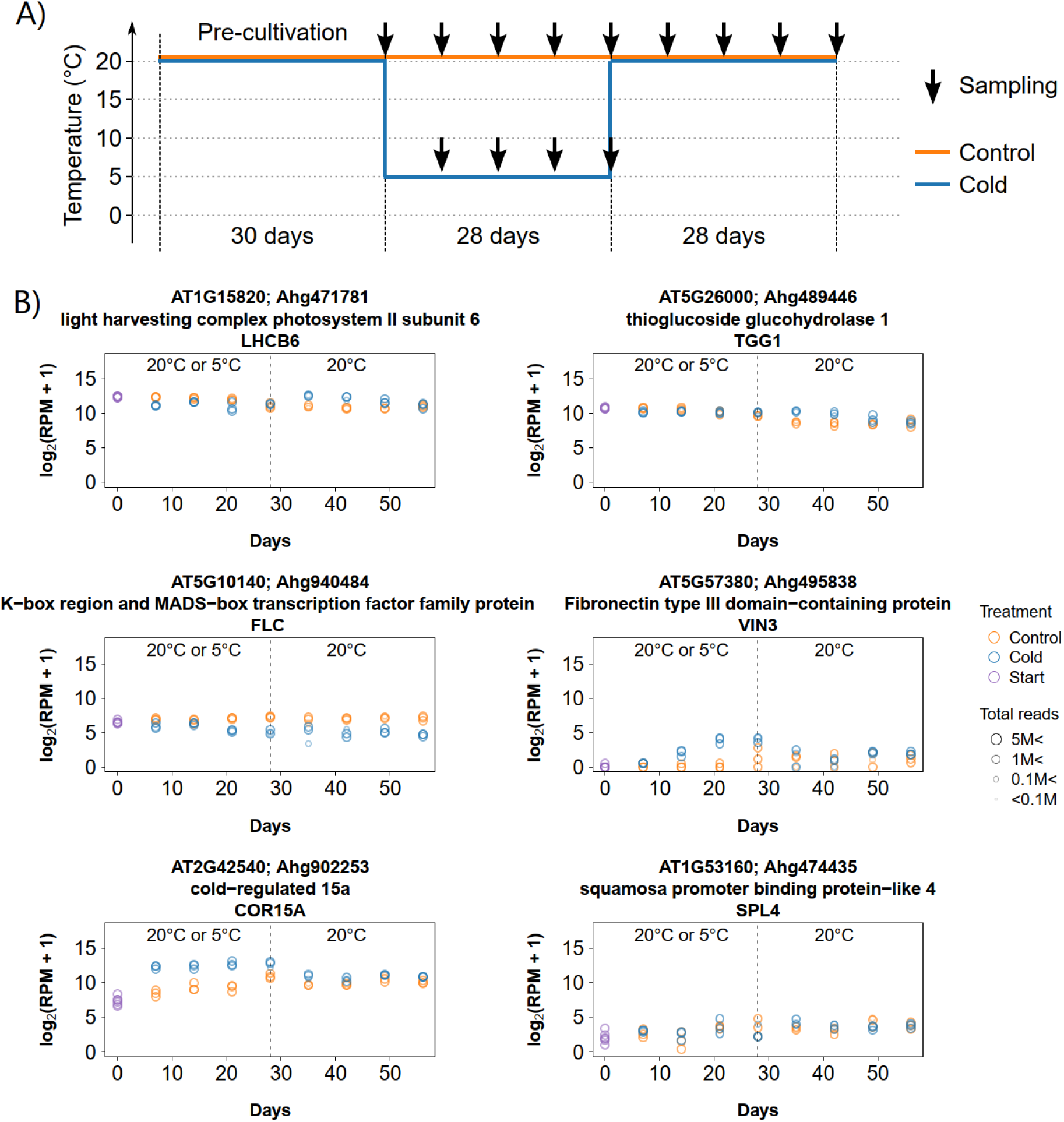
Gene expression in cold exposing experiments against FRI^Sf2^. (A) Experimental scheme. (B) *LHCB6*, *TGG1*, *FLC*, *VIN3*, *COR15A* and *SPL4* are shown. Vertical dashed lines represent the end of cold conditions (4°C).

In the control group, the expression levels of *LHCB6* and *TGG1* decreased as cultivation progressed (Figure 4B). In the cold group, their expression was kept higher than those of the control group after the plants were returned to 20°C. These results showed that the four-weeks cold treatment could not induce the decrease in gene expression of *LHCB6* and *TGG1*. *FLC* expression remained unchanged in the control group, while it continuously downregulated in the cold group. The expression level of *VIN3* remained low in the control group. In contrast, under the cold group, *VIN3* expression increased after three weeks of exposure to 5°C. The expression level then returned to the similar level of the control group one week after the plants were moved back to 20°C. *COR15a* expression increased after one week of exposure to low temperature. These results indicated that the cold treatment worked as expected. No clear difference was observed in the expression level of *SPL4* between the control and cold groups.

## DISCUSSION

Monocarpic plants (mostly annuals) die after flowering, unlike polycarpic plants (mostly perennials). Hence, fitness of monocarpic plants is thought to be maximized by translocating as one’s resources as possible to seed, switching from vegetative phase to reproductive phase in the certain point completely (Kozłowski, 1992). In this study, we surveyed monocarpic *Ath* and polycarpic *Ahg* flowering in spring. Plants contain a high percentage of proteins related to photosynthesis, with about 70-80% of leaf nitrogen residing in chloroplasts (Makino & Osmond, 1991). Cruciferous plants, including *Ath* and *Ahg*, also contain much amount of protein related to glucosinolate metabolism as defense-related proteins against herbivores (Barth & Jander, 2006; Ueda *et al*., 2006; Shirakawa & Hara-Nishimura, 2018). Supporting switching idea, GO term enrichment analysis shows that genes with “autophagy” and “transport” terms are up-regulated only in *Ath*, not in *Ahg*, in spring (February–March) (Figure 3B). Focusing on individual genes, the expression levels of high content proteins which are photosynthesis-related (*LHCB6*) and glucosinolate metabolism genes (*TGG1*) are reduced even before flowering, and the translocation of these resources to seed production is a reasonable response (Figure 2B). To understand the differences in the life histories of *Ath* and *Ahg*, the factors that regulate these genes are needed to clarify. The reduced expression of *LHCB6* and *TGG1* in FRI^Sf2^ from January to April may be also related to developmental aging. *SPL3*, *4* and *5* are genes associated with developmental aging (Jung *et al*., 2011, 2016). Expression level of *SPL4* was saturated in November (Figure 2B), before the decrease in *LHCB6* and *TGG1* expression levels. Low temperatures in winter were considered a candidate environmental factor for producing the unique pattern in gene expression levels of *LHCB6*, and *TGG1* observed in FRI^Sf2^ in spring. The expression of *COR15a* indicate that the low temperature treatment worked well (Figure 4). However, one month of low temperature treatment did not result in both decreased expression of *LHCB6* and *TGG1* and increased expression of *SPL4*. These results show that low temperature is not very important for the down-regulation of *LHCB6* and *TGG1* in spring, and suggest that age may be more important for them. This conclusion is also supported by the previous study (Kurita *et al*., 2021). In the previous study, the expression levels of *LHCB6* and *TGG1* were not reduced in an experiment in which *Ath* was exposed to an environment that mimicked each month for only a few days.

In conclusion, our year-long transcriptome analysis revealed differences in physiological life history strategies between monocarpic *Ath* and polycarpic *Ahg*. Notably, gene expression levels of *LHCB6* and *TGG1* were down-regulated in *Ath* several months before flowering, a pattern not observed in *Ahg*. This suggests that *Ath* initiates the translocation of major proteins (e.g., photosynthesis- and glucosinolate-related proteins) before seed dispersal or mortality to maximize fitness. This process appears to be regulated by age rather than low temperature. However, our study did not capture the full annual transcriptome of *Ath* in its natural habitat, limiting the interpretation of our findings. Nonetheless, the gene expression dynamics related to flowering are likely similar between laboratory and field conditions. This study highlights the significance of long-term transcriptional responses in understanding plant life history. Future comparisons with field-collected annual transcriptome data will further clarify differences in gene expression and reproductive strategies between annual and perennial plants.

## ACKNOWLEDGEMENTS

We thank Kyoko Mogami for technical assistance with the RNA-Seq experiments, and Dynacom Co., Ltd. (Chiba, Japan) for technical assistance with the analysis of RNA-Seq data. This study was supported by JST CREST JPMJCR15O2, JST FOREST JPMJFR210B, JSPS (JP20H00423, JP23H00386, JP23K18156) and MEXT (JP23H04967) for AJN *Arabidopsis* seeds were distributed by ABRC.

## AUTHOR’S CONTRIBUTIONS

AJN conceived the study. YN conducted the experiments and analyzed the data. YN and AJN wrote the manuscript. All authors read and approved the final manuscript.

## DATA AVAILABLITY

The RNA-seq data were submitted to the NCBI Sequence Read Archive repository under the BioProject number PRJNA1223627. R scripts and data required for analysis (count data, etc.) are available via the figshare repository (https://doi.org/10.6084/m9.figshare.30347749)

---

**Figure S1 Bolting date and number of fruits of Col-0 and FRI^Sf2^ under experiments simulating annual cycle.** (A) Bolting date. Tests were conducted using GLMM. Vertical lines represent average of bolting date. (B) In Japan, the annual cycle in temperature lags 1.5 months relative to day length. In the case of *Ahg*, when grown under conditions that eliminate this lag (hereafter in-phase conditions), the number of flowers produced, an indicator of fitness, is lower than when grown under conditions that simulate the 1.5-month lag (hereafter natural conditions), and gene expression patterns are also altered (Nagano *et al*., 2019). To clarify whether the in-phase condition also causes reduced fitness in *Ath*, Col-0 and FRI^Sf2^ were grown under the natural conditions and the in-phase conditions from August to April. Col-0 flowered and died at the end of September (about three months of growth, including pre-culture), so a second experiment was conducted from the end of January (Figure 1B). The bolting date and the number of fruits larger than 5 mm were recorded. The first Col-0 bolted in September-October, the second Col-0 in April, and FRI^Sf2^ in March-April. Col-0 bolted significantly earlier in the in-phase condition than in the natural condition (GLMM, 1st Col-0, *z* = 5.07, *P* = 3.98 × 10^−7^; 2nd Col-0, *z* =2.7, *P* = 6.07 × 10^−3^) (Figure S1). FRI^Sf2^ tended to bolt earlier under in-phase conditions, but not significantly (GLMM, *z* = 1.63, *P* = 0.103). Interaction of bolting date and the conditions (natural and in-phase) on fruit numbers was significant for both Col-0 and FRI^Sf2^ (GLMM, 1st Col-0, *χ*^2^ = 13.5, *P* = 2.41× 10^−4^; 2nd Col-0, *χ*^2^ = 3.9, *P* = 0.0482; FRI^Sf2^, *χ*^2^ = 11.7, *P* = 6.33×10^−4^). In other words, the effect of the bolting date on the number of fruits differed depending on the conditions. In all cases, under the natural conditions, the bolting date had little effect on the number of fruits, while under the in-phase conditions, the later the bolting date, the fewer the number of fruits. This suggests that under the natural conditions, fruit numbers were robust to variations in the bolting date, while under “unnatural” conditions such as the in-phase, the later the bolting date, the less fitness than that under the natural conditions. In this study, the number of fruits was significantly higher in the in-phase conditions than in the natural conditions in the first Col-0 (GLMM, 1st Col-0, *z* = 2.13, *P* = 0.0334). This may be due to the fact that the first Col-0 bolted early. There were no significant differences in fruit number among the remaining conditions (GLMM, 2nd Col-0, *χ*^2^ = −0.91 *P* = 0.364 FRI^Sf2^, *χ*^2^ = 1.01, *P* = 0.311). These results differ from those of previous studies of *A. halleri* (Nagano *et al*., 2019), the reasons for which are unknown.

**Figure S2 Summary of sample sets and gene sets in this study.** Samples analysed by RNA-Seq in this study and samples above the read number filter (> 10^5.5^ reads). The histogram of total read numbers is shown in the inset. The dashed line indicates the threshold of 10^5.5^ reads. Gene sets analysed in this study. The histogram of mean expressions is shown in the inset.

**Figure S3 Pearson correlation between the monthly average of each transcript profile including expanded data of AthGCShort by 1-, 3- and 7-days treatments**. Correlation coefficients were calculated using the monthly average of log-transformed RPMs for the variable genes (n = 12,141 genes).

**Table S1 Sample description of RNA-Seq in GC experiment simulating annual cycle**

**Table S2 Setting for first Col-0 and FRI^Sf2^ GC experiment simulating annual cycle**

**Table S3 Setting for second Col-0 GC experiment simulating annual cycle**

**Table S4 Sample description of RNA-Seq in GC experiment exposing cold**

**Table S5 Setting for FRI^Sf2^ GC experiment exposing cold**

**Table S6 GO enrichment analysis of DEGs in Feb-Mar not in Oct-Nov**

