## Supplemental Figures S1-S3 for "Year-long transcriptome survey highlights a resource allocation strategy at the late growth stage in a monocarpic plant"

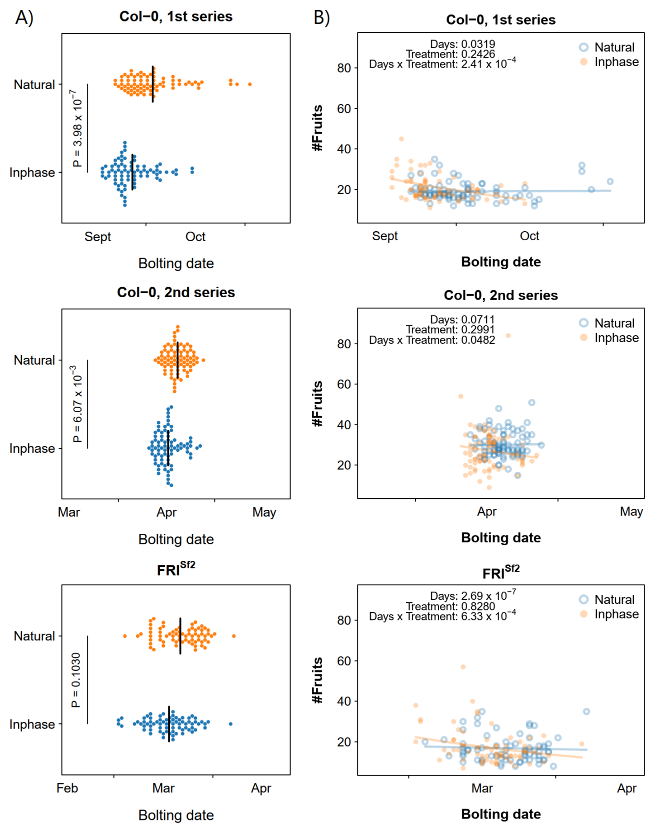


**Figure S1 Bolting date and number of fruits of Col-0 and FRISf2 under experiments simulating annual cycle.** (A) Bolting date. Tests were conducted using GLMM. Vertical lines represent average of bolting date. (B) In Japan, the annual cycle in temperature lags 1.5 months relative to day length. In the case of Ahg, when grown under conditions that eliminate this lag (hereafter in-phase conditions), the number of flowers produced, an indicator of fitness, is lower than when grown under conditions that simulate the 1.5-month lag (hereafter natural conditions), and gene expression patterns are also altered (Nagano et al., 2019). To clarify whether the in-phase condition also causes reduced fitness in Ath, Col-0 and FRISf2 were grown under the natural conditions and the in-phase conditions from August to April. Col-0 flowered and died at the end of September (about three months of growth, including pre-culture), so a second experiment was conducted from the end of January (Figure 1B). The bolting date and the number of fruits larger than 5 mm were recorded.

The first Col-0 bolted in September-October, the second Col-0 in April, and FRISf2 in March-April. Col-0 bolted significantly earlier in the in-phase condition than in the natural condition (GLMM, 1st Col-0, z = 5.07, P = 3.98 × 10-7; 2nd Col-0, z =2.7, P = 6.07 × 10-3) (Figure S1). FRISf2 tended to bolt earlier under in-phase conditions, but not significantly (GLMM, z = 1.63, P = 0.103). Interaction of bolting date and the conditions (natural and in-phase) on fruit numbers was significant for both Col-0 and FRISf2 (GLMM, 1st Col-0, $\chi 2$ = 13.5, P = 2.41 × 10-4; 2nd Col-0, $\chi 2$ = 3.9, P = 0.0482; FRISf2, $\chi 2$ = 11.7, P = 6.33 × 10-4). In other words, the effect of the bolting date on the number of fruits differed depending on the conditions. In all cases, under the natural conditions, the bolting date had little effect on the number of fruits, while under the in-phase conditions, the later the bolting date, the fewer the number of fruits. This suggests that under the natural conditions, fruit numbers were robust to variations in the bolting date, while under "unnatural" conditions such as the in-phase, the later the bolting date, the less fitness than that under the natural conditions. In this study, the number of fruits was significantly higher in the in-phase conditions than in the natural conditions in the first Col-0 (GLMM, 1st Col-0, z = 2.13, P = 0.0334). This may be due to the fact that the first Col-0 bolted early. There were no significant differences in fruit number among the remaining conditions (GLMM, 2nd Col-0, $\chi 2$ = -0.91 P = 0.364 FRISf2, $\chi 2$ = 1.01, P = 0.311). These results differ from those of previous studies of A. halleri (Nagano et al., 2019), the reasons for which are unknown.

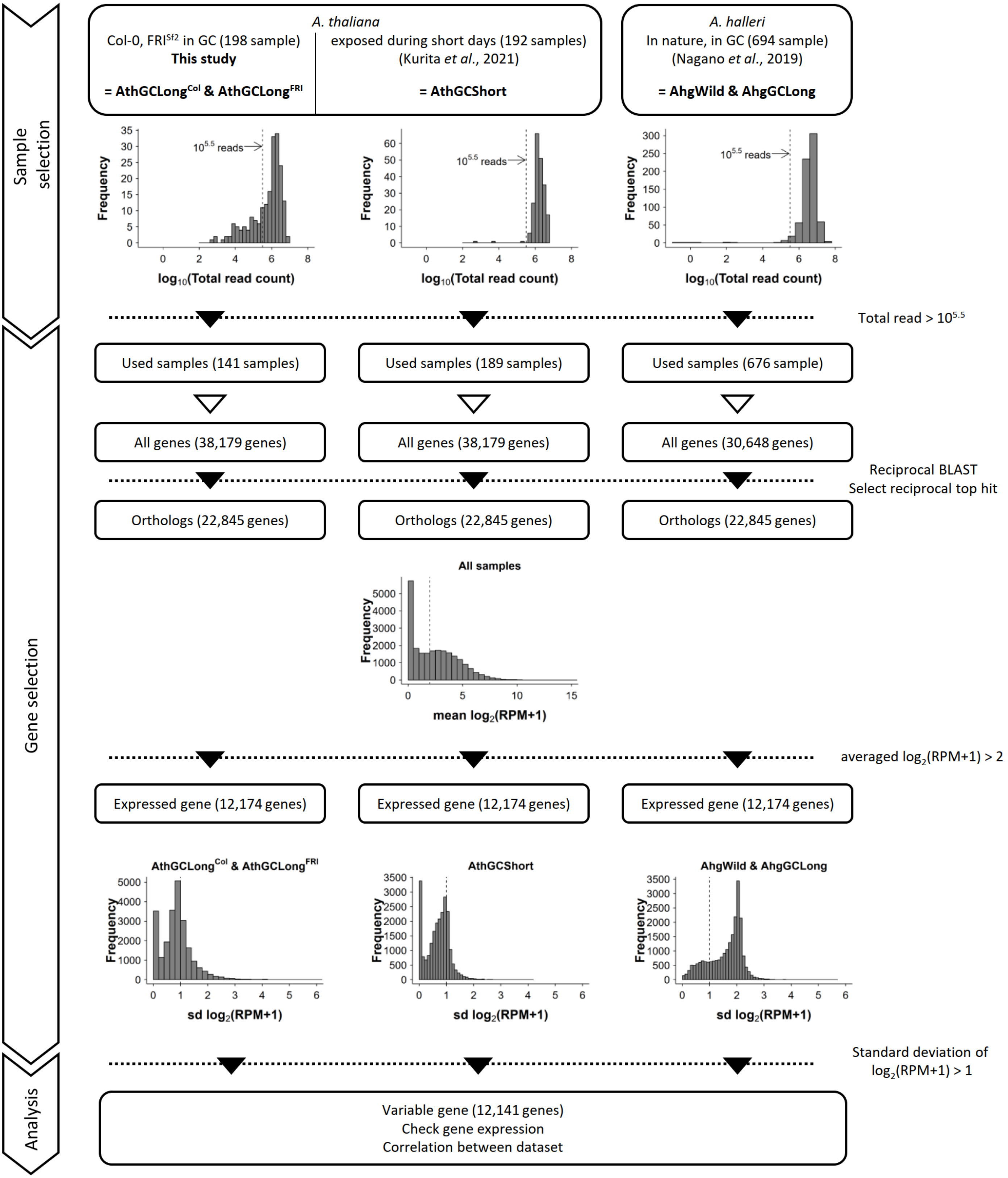


**Figure S2 Summary of sample sets and gene sets in this study.** Samples analysed by RNA-Seq in this study and samples above the read number filter (> $10^{5.5}$ reads). The histogram of total read numbers is shown in the inset. The dashed line indicates the threshold of $10^{5.5}$ reads. Gene sets analysed in this study. The histogram of mean expressions is shown in the inset.

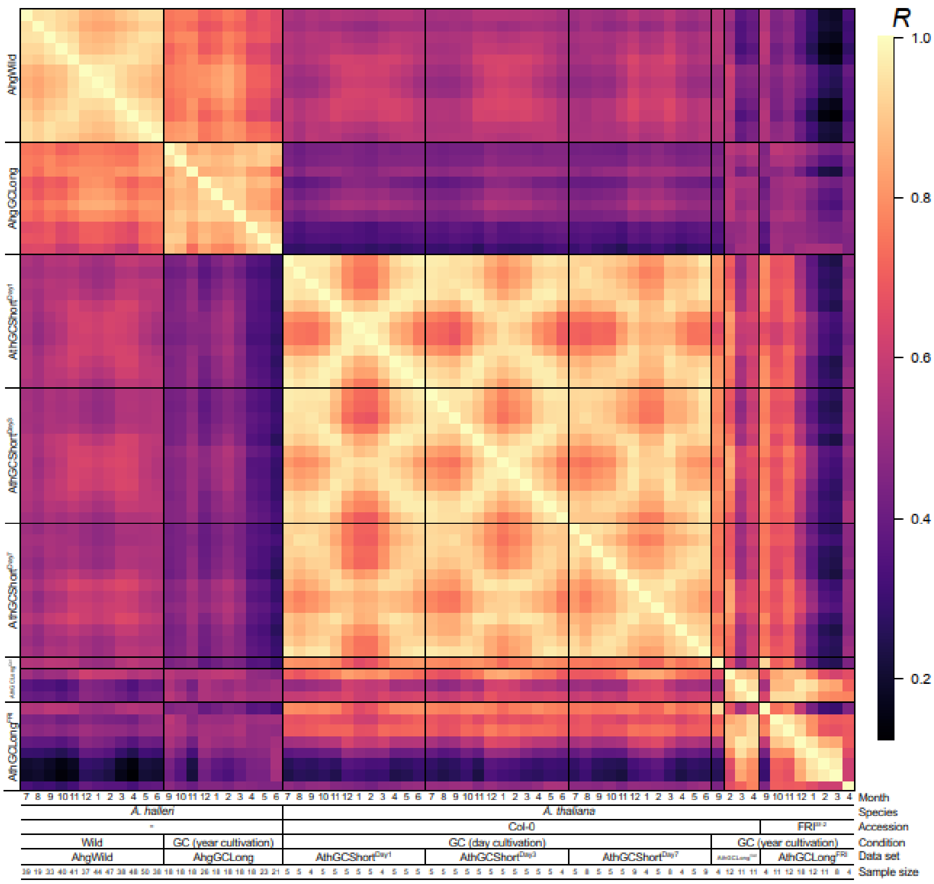


**Figure S3 Pearson correlation between the monthly average of each transcript profile including expanded data of AthGCShort by 1-, 3- and 7-days treatments.** Correlation coefficients were calculated using the monthly average of log-transformed RPMs for the variable genes (n = 12,141 genes).
